# Reduced responsiveness of the reward system underlies tolerance to cannabis impairment in chronic users

**DOI:** 10.1101/708677

**Authors:** N.L Mason, E.L. Theunissen, N.R.P.W. Hutten, D.H.Y. Tse, S.W. Toennes, J.F.A. Jansen, P. Stiers, J.G. Ramaekers

## Abstract

Cannabis is the most commonly used illicit drug in the world. However due to a changing legal landscape, and rising interest in therapeutic utility, there is an increasing trend in (long-term) use and possibly, cannabis impairment. Importantly, a growing body of evidence suggests regular cannabis users develop tolerance to the impairing, as well as the rewarding, effects of the drug. However, the neuroadaptations that may underlie cannabis tolerance remain unclear. Therefore, this double-blind, randomized, placebo controlled, cross-over study assessed the acute influence of cannabis on brain and behavioral outcomes in two distinct cannabis user groups. Twelve occasional (OUs) and 12 chronic (CUs) cannabis users received acute doses of cannabis (300 μg/kg THC) and placebo, and underwent ultra-high field functional magnetic resonance imaging (fMRI) and magnetic resonance spectroscopy (MRS). In OUs, cannabis induced significant neurometabolic alterations in reward circuitry, namely decrements in functional connectivity and increments in striatal glutamate concentrations, which were associated with increases in subjective high and decreases in performance on a sustained attention task. Such changes were absent in CUs. The finding that cannabis altered circuitry and distorted behavior in OUs, but not CUs, suggests reduced responsiveness of the reward circuitry to cannabis intoxication in chronic users Taken together, the results suggest a pharmacodynamic mechanism for the development of tolerance to cannabis impairment.

## Introduction

Cannabis is the most commonly used illicit drug in the world, with 4% of the global population reportedly using the substance (1). However due to a changing legal landscape, and rising interest in therapeutic utility, there is an increasing trend in (long-term) use (1, 2). Importantly, a growing body of evidence suggests that the acute effects of cannabis are less prominent in regular cannabis users (3), suggesting development of tolerance to the impairing, as well as the rewarding, effects of the drug. Nonetheless, the neurobiological mechanisms underlying cannabis tolerance are unknown.

Accumulating evidence suggests that the main psychoactive component of cannabis [delta-9-tetrahydrocannabinol (THC)] binds to cannabinoid (CB1) receptors located on GABAergic and glutamatergic neurons distributed throughout the brain, with high densities found in limbic-reward structures (4). Subsequently, THC has been found to acutely activate the reward circuitry, increasing dopamine (5–9) and glutamate (10, 11) concentration levels in key brain areas including the striatum, nucleus accumbens (NAc) and prefrontal cortex (PFC); a pattern implicated in both the rewarding and impairing effects of drugs of abuse (12, 13).

Accordingly, studies with *chronic* cannabis users have found alterations in dopaminergic function in striatal areas (14–16), as well as decreases in glutamate concentrations in the basal ganglia (17, 18) and anterior cingulate cortex (ACC) (19, 20). Furthermore, repeated use of cannabis has been associated with structural changes in frontal areas, as evinced by decreased grey matter volume (21–23), and abnormal concentrations of metabolites including N-acetylaspartate (NAA), myo-inositol (mI), and choline containing compounds (Cho), biochemical markers of neuronal integrity and glial activation (18, 19, 24–28). Taken together, these studies provide evidence that repeated cannabis exposure may lead to alterations in neurotransmission and neuronal health, which underlie the diminished cognitive and behavioral response associated with acute cannabis tolerance. However, to date the neuroadaptations that may underlie cannabis tolerance have not been systematically assessed.

Therefore the aim of the present double-blind, placebo controlled study was twofold. The first goal was to assess acute influence of cannabis in two different cannabis using groups, namely occasional (OUs) and chronic (CUs) users, on brain and behavioral outcomes previously found to be affected by cannabis. To do this, both OUs and CUs received a single cannabis dose containing 300 μg/kg THC. Ultra-High Field (7T) proton magnetic resonance spectroscopy (^1^H MRS), a non-invasive imaging technique that allows reliable in vivo measurement of neurometabolites and neurotransmitters, was used to assess glutamate, gamma-aminobutyric acid (GABA), NAA, Cho, and mI levels in the striatum and ACC. Resting state functional magnetic resonance imaging (fMRI) data were acquired to determine functional connectivity between the regions of interest (ROI) in the NAc and remote cortical areas, as an indirect measure of dopaminergic stimulation (10, 29, 30). Furthermore *a priori* ROI-to-ROI analysis assessed differences in connectivity strength between areas of the reward circuit. Finally, subjective high and sustained attention, two outcome variables shown to be modulated by (acute) cannabis exposure (10, 31, 32), were assessed.

The second goal was exploratory in nature, to evaluate long-term effects of repeated cannabis exposure, by comparing the placebo condition of each group. Overall, we hypothesized that THC would induce behavioral, functional, and metabolic changes in the OUs, but not the CUs, indicative of (neuroadaptive) tolerance. Furthermore, based on previous studies with chronic cannabis users, we hypothesized that during placebo, CUs would show decreased concentrations of metabolites compared to OUs.

## Methods

### Participants

In total, 27 healthy cannabis users entered the study. Three participants were excluded due to poor fMRI data, resulting in a total of 12 occasional and 12 chronic users (male N= 14, female N= 10). For further details see SI Methods.

This study was conducted according to the code of ethics on human experimentation established by the declaration of Helsinki (1964) and amended in Fortaleza (Brazil, October 2013) and in accordance with the Medical Research Involving Human Subjects Act (WMO) and was approved by the Academic Hospital and University’s Medical Ethics committee. All participants were fully informed of all procedures, possible adverse reactions, legal rights and responsibilities, expected benefits, and their right for voluntary termination without consequences.

### Design, doses, and administration

This study was conducted according to a double-blind, placebo-controlled, mixed cross-over design. Treatments consisted of placebo and 300 μg/kg THC (Bedrobinol; 13.5 % THC) on separate days, separated by a minimum wash-out period of 7 days for OUs, to avoid cross-condition contamination. Treatment orders and doses were randomly assigned to subjects according to a balanced, block design. The dosage of cannabis was tailored to individual participants in order to reach 300 μg/kg bodyweight THC, which has previously been found to be well tolerated by subjects with an average experience of cannabis use (33, 34). For an overview of the testing day schedule, see Table S5.

### Proton MR Spectra

Single-voxel proton magnetic resonance spectroscopy (MRS) measurements were performed on a MAGNETOM 7T MR scanner (Siemens Healthineers, Erlangen, Germany) with a whole-body gradient set (SC72; maximum amplitude, 70 mT/m; maximum slew rate, 200 T/m/s) and using a single-channel transmit/32-channel receive head coil (Nova Medical, Wilmington, MA, USA). Spectroscopic voxels of interest were placed by a trained operator at the Anterior Cingulate Cortex (ACC) (voxel size = 25 × 20 × 17 mm^3^) and the right striatum (voxel size = 20 × 20 × 20 mm^3^). Spectra were acquired with stimulated echo acquisition mode (STEAM) (35) sequence using the following parameters: TE = 6.0 ms, TM = 10.0 ms, TR = 5.0 s, NA = 64, flip angle = 90°, RF bandwidth = 4.69 kHz, RF centred at 2.4 ppm, receive bandwidth = 4.0 kHz, vector size = 2048, 16-step phase cycling, acquisition time = 5:20 min. Outcome measures for MRS were concentration ratios of glutamate, GABA, NAA, mI, and Cho to total Creatine (tCr, Creatine + Phospho-Creatine). For further information and fit quality measures see SI Methods and Figure S2 and Table S6.

### Resting state fMRI

During resting state, 258 whole-brain EPI volumes were acquired (TR=1400 ms; TE= 21 ms; field of view (FOV)=198 mm; flip angle=60°; oblique acquisition orientation; interleaved slice acquisition; 72 slices; slice thickness=1.5 mm; voxel size=1.5×1.5×1.5 mm). During resting state scans, participants were shown a black cross on a white background, and were instructed to focus on the cross while attempting to clear their mind, and lay as still as possible. For further details see SI Methods.

### Functional Connectivity

Functional connectivity data were produced with the MATLAB toolbox DPARSF (36). In order to indirectly assess dopamine neurotransmission, two spheres (4 mm radius) were created that were located (in MNI space) in the left and right NAc. Average time courses were obtained for each sphere separately and correlational analysis was performed voxel wise to generate functional connectivity maps for each sphere.

Furthermore, as we were interested in FC *within* the reward reward circuit, ROI-to-ROI FC was computed according to the same aforementioned procedure, between areas including: NAc, MDN, VPN, and MC. Fisher’s correlation coefficient maps were created between the NAc and MDN, NAc and VPN, MDN and VPN, MDN and MC, and MCand NAc. For further details see SI Methods.

### Psychomotor Vigilance Task

The psychomotor vigilance task (PVT) is a sustained-attention, reaction-time task that measures the speed with which participant respond to a visual stimulus (37). The participant is instructed to press a button as soon as the stimulus appears (red circle). The outcome measures of the task are response speed (mean reaction time) and number of attentional lapses (reaction time > 500 ms). For more information see SI Methods.

### Subjective High

Participants rated their subjective high on visual analogue scales (10 cm) on four consecutive time points after treatment administration. Participants had to indicate how high they felt at that moment, compared with the most high they have ever felt (0=not high at all; 10=extremely high).

### Pharmacokinetic Measures

Blood samples (8 mL) to determine cannabinoid concentrations (THC and metabolites OH-THC and THC –COOH) were taken at base- line, 10, 30, 50, and 70 minutes post administration. Blood samples were centrifuged and serum was frozen at −20 °C until analyses for pharmacokinetic assessments. Cannabinoid concentrations were determined using a validated and proficiency test approved forensic routine method consisting of an automated solid-phase extraction and gas chromotrography with tandem mass spectrometric detection with a limit of quantification of 0.3 ng/ml or less (10).

#### Statistical Analysis

##### Subjective high, sustained attention, and metabolite concentrations

Statistical analysis of subjective high, task performance, and metabolite concentrations were conducted in IBM SPSS Statistics 24 using a mixed model analysis consisting of the within-subject factors treatment (THC and placebo), and time after smoking (2 levels), and the between-subject factor of group (OU or CU). Due to main effect of Treatment or interaction of Treatment X Group, a second analysis was performed for each group, with treatment and time as within-subject factors. The alpha criterion level of significance was set at p=0.05. Due to a violation of the assumption of normality, the data for the number of lapses and mean reaction time were log transformed.

##### fMRI data

Functional connectivity data (i.e. correlation coefficient maps for each individual in each treatment condition at each time point) were analyzed in a GLM model in SPM 12.

For the reward circuit ROI-to-ROI analysis of Fisher’s correlation coefficient values was conducted in IBM SPSS statistics 24. For each group, a repeated measures analysis was conducted consisting of the within-subject factor treatment (THC and placebo). For details, see SI Methods.

##### Correlation analysis

Correlation analyses were conducted to further investigate the relationship between cannabis induced changes in brain and behavior. Correlation input included average treatment change values of *(i)* behavioral outcomes (subjective high and number of lapses), *(ii)* MRS concentration levels, *(iii)* ROI-to-ROI Fisher’s correlation values, and *(iv)* mean voxel activation of SPM identified clusters from a voxel wise correlation analysis between NAc FC and behavioral outcomes. Pearson’s correlations were performed in SPSS. For details, see SI Methods. For overview of investigated correlations, see Table S7.

##### Exploratory analysis

Separate analyses of covariance (ANCOVAS) were carried out for average (placebo) neurometabolite concentrations, behavioral measures, and FC in the reward circuit, as the dependent variables and group (OU vs CU) as the fixed factor. Serum THC, 11-OH-THC, and THC-COOH levels were entered as covariates, due to significant differences between groups.

## Results

### Demographic Characteristics

OUs (n = 12) and CUs (n = 12) did not differ with respect to gender distribution, age, history of cannabis use, or consumption of alcohol, caffeine, nicotine, or other drugs (Table S1). As expected, CUs reported using significantly more cannabis per week than OUs.

### THC Concentrations in Serum

Mean (SE) concentrations of THC, 11-OH-THC, and THC-COOH in serum are given in Table S2. As expected from previous experience (38), CUs not only exhibited significantly higher THC-COOH concentrations, but also reached significantly higher THC levels from the same dose regimen than OUs.

### Subjective and Cognitive Effects

Mixed model ANOVA [Treatment(THC vs placebo) *Time(timepoint 1 vs timepoint 2)*Group(OUs vs CUs)] yielded a significant main effect of Treatment [F(1,21) = 49.682, P< .0001, ηp²=.703] and Time [F(3,63) = 42.269, P< .0001, ηp²=.668], and a significant interaction of Treatment*Group [F(1,21)=5.023, P=.036, ηp²=.193] and Treatment*Time [F(363) = 17.346, P< .0001, ηp²=.452] on subjective high, indicating that THC increased feelings of subjective high in both groups, but to a larger degree in OUs (Figure S1A).

Analysis yielded a significant main effect of Treatment [F(1,20) = 8.057, P= .010, ηp²=.287] and Time [F(1,20) = 7.620, P=.012, ηp²=.276], and a significant interaction of Time*Group[F(1,20) = 8.085, P=.010, ηp²=.288] on mean reaction time of the PVT. Further analysis revealed that mean reaction time was significantly increased by THC in OUs [F(1,10) = 5.226, P= .045, ηp²=.343], but not in CUs [P>.1] (Figure S1B).

Analysis yielded a significant main effect of Treatment [F(1,21) = 7.793, P= .011, ηp²=.271], and a significant interaction of Time*Group[F(1,21) = 4.313, P= .050, ηp²=.170] on number of attentional lapses during the PVT. Further analysis revealed number of attentional lapses was significantly increased by THC in OUs [F(1,10) = 5.286, P= .044, ηp²=.346], but not in CUs [P>.1] (Figure S1C).

#### Metabolite Concentrations

##### Striatum

Mixed model ANOVA yielded a significant interaction of Treatment *Time *Group [*F*(1,21)=8.779, *P*=.007, ηp²=.295] on glutamate/tCr (Glu) concentration levels. Further analysis revealed that concentration levels were significantly increased by THC in OUs [*F*(1,11) = 11.506, *P*= .006, ηp²=.511], but not in CUs [*P*>.1] (Figure 1*A*)].

**Figure 1.**
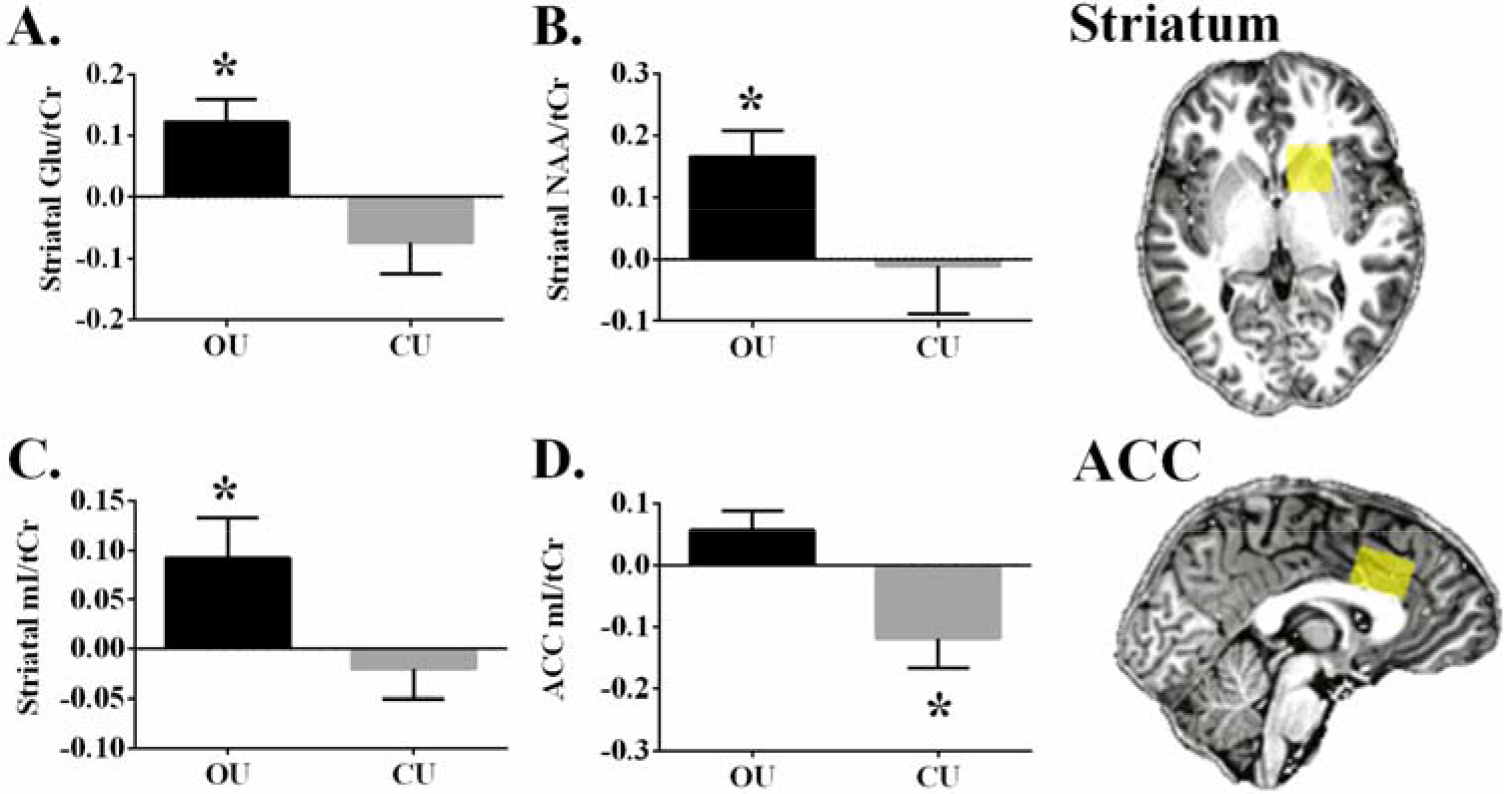
Occasional and chronic users mean (SE) metabolite concentration levels, averaged over both timepoints for both treatments [average(THC timepoint 1 – Placebo timepoint 2; THC timepoint 2 – Placebo timepoint 2)] A. Striatal glutamate. B. Striatal total n-acetyl-aspartate. C. Striatal myoinositol. D. Anterior cingulate cortex myoinositol. *within group analysis, P<.05

Analysis yielded a significant interaction of Treatment *Time *Group [*F*(1,19)=6.546, *P*=.019, ηp²=.256] and a significant main effect of Treatment [*F*(1,19) = 4.670, *P*=.044, ηp²=.197] on NAA + NAAG/tCr (NAA) concentration levels. Further analysis revealed that concentration levels were significantly increased by THC in OUs [*F*(1,11) = 15.345, *P*= .002, ηp²=.582], but not in CUs [*P*>.1] (Figure 1*B*).

Analysis yielded a trending interaction of Treatment*Group [*F*(1,20)=4.241, *P*=.053, ηp²=.175] on mI/tCr (mI) concentration levels. Further analysis revealed that concentration levels were significantly increased by THC in OUs [*F*(1,11) = 5.176, *P*= .044, ηp²=.320], but not in the CUs [*P*>.1] (Figure 1*C*).

##### Anterior cingulate cortex

Mixed model ANOVA yielded a significant interaction of Treatment*Group [*F*(1,18)=7.450, *P*=.014, ηp²=.293] on mI concentration levels. Further analysis revealed that concentration levels were significantly decreased by THC in CUs [*F*(1,9) = 5.935, *P*= .038, ηp²=.397], but not in OUs [*P*>.1] (Figure 1*D*)].

Other metabolites did not reach significance (see Table S3 for mean metabolite concentrations).

##### Functional Connectivity as an Indirect Measure of Dopamine

The contrast drug vs placebo (Placebo > THC) resulted in reduced functional connectivity with the NAc seeds in both hemispheres in OUs, whereas no change was found in CUs (Figure 2). Reductions in functional connectivity were prominent in broad areas of the frontal, temporal, parietal, and occipital lobes, a pattern typical of an increase in dopaminergic neurotransmission (Table S4). No significant differences in activation were found for the inverse comparison (THC > Placebo) in either group. Furthermore, no significant differences were seen between the left and right NAc seed, so only left NAc seed results are shown.

**Figure 2.**
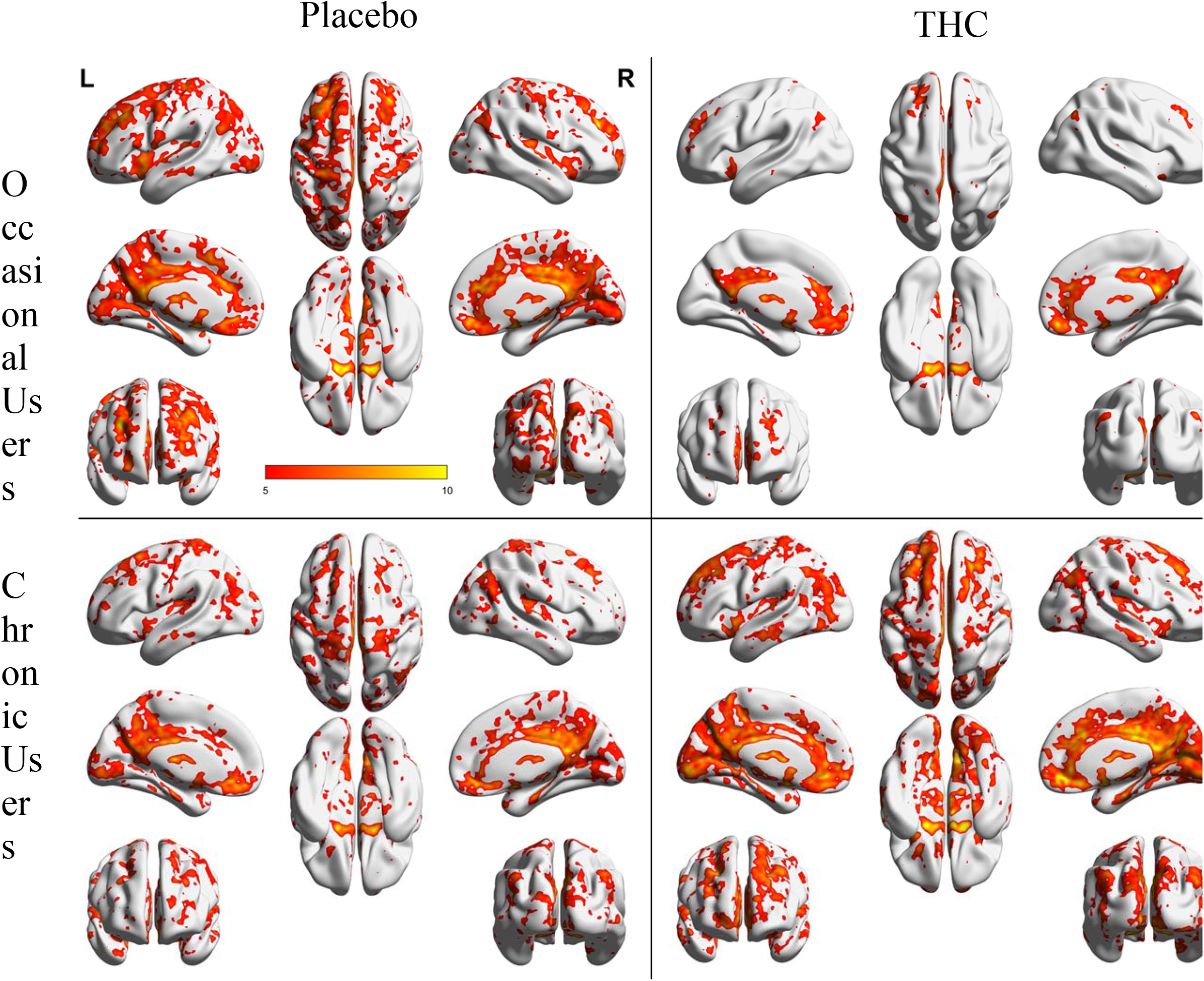
NAcc-related functional connectivity in the left hemispheres. Shown are thresholded Z-score maps of functional connectivity for each group, and each condition.

##### Functional Connectivity in the Reward Circuit

Results of the region-of-interest (ROI)-to-ROI functional connectivity analysis are displayed in (Figure 3). ROIs were chosen because they are established structures of the reward circuitry, a cortico-subcortical network connected via glutamatergic and GABAergic projections between the NAc, ventral pallidum (VP), medial dorsal nucleus (MDN), and the prefrontal cortex (39–42) (Figure 3). NAc, VP, and MDN seeds were chosen *a priori*, whereas the prefrontal seed was chosen based on the previous FC analysis, which indicated significant treatment (Placebo > THC) induced changes in the midcingulate area (MC). Mixed model ANOVA revealed that cannabis decreased functional connectivity between the ROIs in OUs, whereas no significant treatment effect was seen in CUs.

**Fig. 3.**
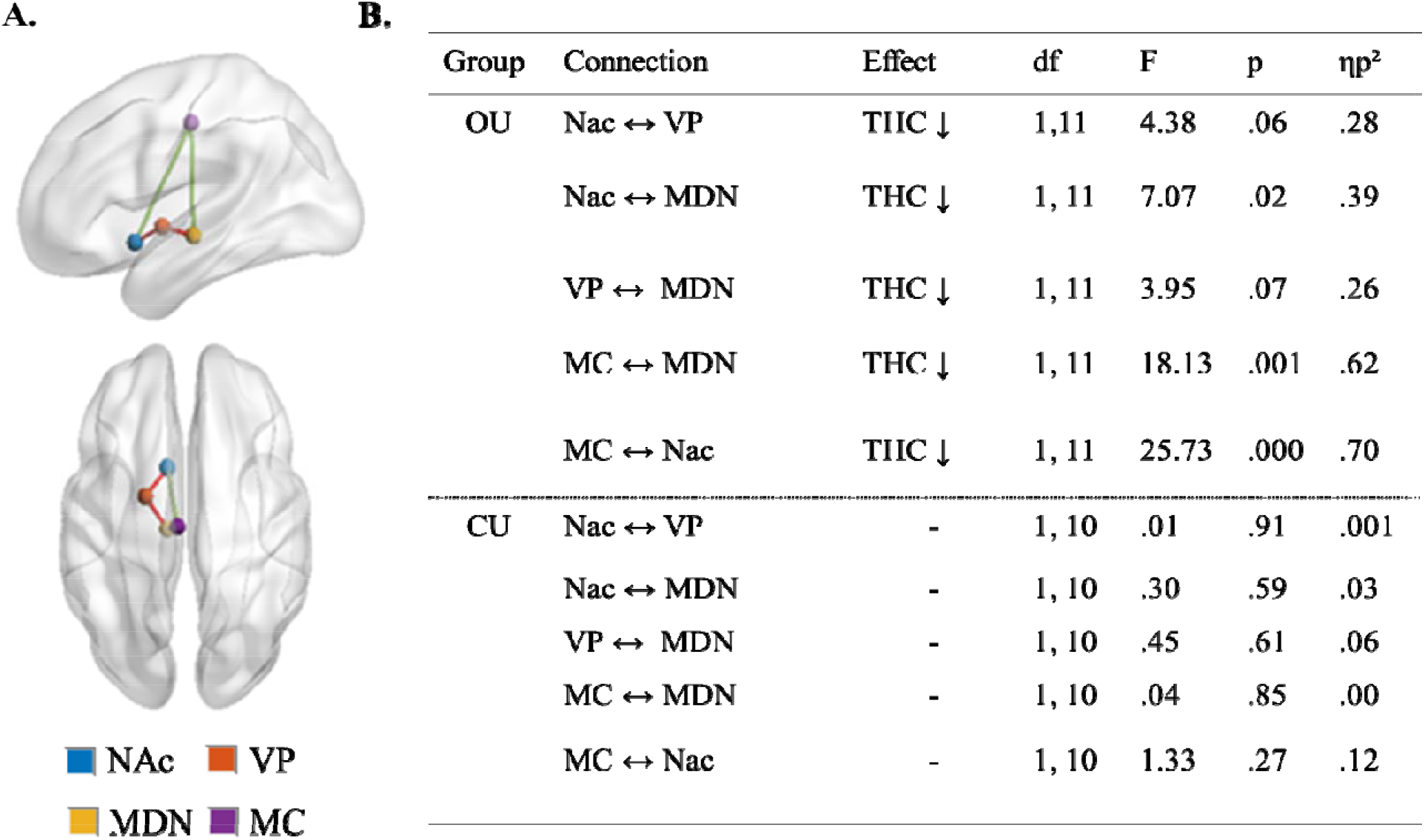
ROI-to-ROI FC analysis results. A) Four nodes of the reward circuit, where ROIs were placed. Red lines pertain to gabaergic pathways, whereas green lines pertain to glutamatergic pathwa s. NAc = nucleus accumbens; VP = ventral pallidum; MDN = medial dorsal nucleus; MC = midcingulate area B) Results of repeated measures analysis (THC vs placebo), separated by group.

##### Relationship Between cannabis Induced Changes in Brain and Behavior

Correlation analyses were conducted to evaluate the association between treatment induced changes in (*i*) subjective high and performance on the sustained attention task, (*ii*) Glu, NAA, and mI concentration levels, and (*iii*) FC in the reward circuit. Analysis revealed a significant positive correlation in OUs (r=.641) between treatment induced changes in striatal NAA and number of attentional lapses, and a significant negative correlation (r=−.618) between treatment induced changes in striatal mI and subjective high. For an overview of all correlational analyses, see Table S7. A further voxel wise correlation analysis between NAc FC whole brain correlation coefficient maps and behavioral outcomes was performed, revealing significant positive correlations (all r > .6) in FC between the NAc and cortical brain areas, and behavioral outcomes (Table S8).

##### Exploratory Analysis

In order to assess long-term effects of repeated cannabis exposure, metabolite concentrations, reward circuit FC, and behavioral outcomes during the placebo condition of each group, were compared. The analysis was exploratory in nature, due to the low sample size. No significant difference between groups was found in any of the variables of interest, when controlling for THC and THC-metabolite concentrations in blood (*P*>.1).

## Discussion

The present study demonstrates the first attempt to assess cannabis induced neuroadaptations in the reward system, which may underlie behavioral cannabis tolerance. Using an ultra-high field multimodal brain imaging approach, we showed that in occasional users, cannabis induced significant neurometabolic alterations in the reward circuitry, namely decrements in FC and increments in striatal glutamate, which were associated with increases in subjective high and decreases in performance on a sustained attention task. Such changes were absent in chronic cannabis users. The finding that cannabis altered reward circuitry and distorted behavior in OUs, but not CUs, suggests the development of neuroadaptations in the reward circuitry after excessive use of cannabis that reduce the circuitry and behavioral response to acute cannabis impairment.

The present study found that, in OUs, cannabis decreased coupling between BOLD responses in keys areas of the reward system, a fronto-subcortical network of brain structures that are connected via dopaminergic, GABAergic, and glutamatergic neurotransmission (i.e. nucleus accumbens, VTA, ventral pallidum), motivation and salience attribution (i.e. medial orbital frontal cortex), executive and inhibitory control (i.e. anterior cingulate cortex) and conditioning and memory (i.e. amygdala, medial orbital frontal cortex, hippocampus) (43, 44). Importantly, decrements in FC between the NAc and regions such as the thalamus and frontal cortex have been suggested to reflect *increase*s in dopaminergic neurotransmission throughout the circuit (10, 29, 30). Specifically, the NAc receives dopaminergic input from the VTA, which is under inhibitory control of GABA interneurons on which presynaptic CB1 receptors (CB1R) are located. Stimulation of CB1R by THC disinhibits the VTA, which in turn increases dopamine levels in the NAc (45). Subsequently, the increase in striatal dopamine level output from the NAc decreases the GABAergic inhibitory tone to the thalamus, which is reflected in decreased FC, as seen in this study.

Furthermore, this process of disinhibition has been suggested to lead to increased glutamatergic signaling to the prefrontal cortex, subsequently to the VTA, and back to the NAc (30). Accordingly, MRS in OUs showed cannabis increased striatal glutamate concentration levels; a finding that is compatible with previous human studies which have found acute increases (10, 11), as well as chronic decreases (18) of striatal glutamate after exposure to THC. Furthermore, it has previously been demonstrated that cannabis induced changes in striatal glutamate levels correlate strongly to cannabis induced alterations of functional connectivity within the fronto-subcortical circuit (10). Thus it could be hypothesized that stimulatory glutamatergic input from the prefrontal cortex to the NAc synergizes with cannabis induced increases in dopaminergic input from the VTA to the NAc, and further strengthens the disinhibition of thalamic signaling in the fronto-subcortical circuitry. However, MRS did not show altered ACC glutamate concentrations in the OU group, as would be hypothesized. Indeed previous results regarding the effects of cannabis on glutamate in the ACC have been mixed, with studies by one group finding reductions of glutamate-related metabolite concentrations in cannabis users (19, 20), whereas no alterations were found in a subsequent chronic (28), or acute (10) study. Nonetheless, previous work utilizing microdialysis found THC increased extracellular glutamate in the rat prefrontal cortex (9), supporting the hypothesis that THC disrupts glutamatergic signaling in frontal areas, however this effect may not be localized to the ACC. Thus future studies should assess the impact THC has on other areas in the frontal cortex, like the medial orbitofrontal cortex.

The present study also demonstrated that cannabis increased subjective high and decreased performance on a sustained attention task in OUs, outcome variables previously found to be affected by cannabis (10, 31, 32). Furthermore, behavioral outcomes correlated with striatal functional connectivity to other areas of the reward circuit. Accordingly, cannabis induced changes in striatal glutamate and striatal FC have been significantly associated with decrements in cognitive function and impulse control (30), as well as increases in subjective high (10, 30) and experience of psychotomimetic symptoms (11). Taken together, the findings suggest that the impact of cannabis on neural activity within the reward circuit may underlie multiple behavioral changes observed after acute cannabis exposure.

In line with this, the CUs demonstrated an absence of cannabis induced stimulation of the reward circuit, as well as mitigation of behavioral alterations. Specifically, no changes were seen in either FC between areas of the reward circuitry, or glutamate concentration levels, when comparing cannabis to placebo. Furthermore, sustained attention performance did not significantly differ between treatment conditions. However, CUs reported significantly increased levels of subjective high after cannabis relative to placebo, although the change in high was to a lesser extent than in the OUs, as expected (31). Taken together, findings suggest that chronic cannabis users exhibit pharmacodynamic tolerance to the effects of cannabis.

The mechanisms by which neurobiological tolerance to the acute effects of THC develops have yet to be fully elucidated. Animal and human research generally supports the notion of CB1R downregulation and desensitization in cortical and subcortical regions after repeated exposure to cannabis (46–53). Although studies have reported global reduction in CB1R availability in chronic cannabis users, it has been suggested that neuroadaptive changes take place in a time and region-specific manner (54), with regional analysis demonstrating significant CB1R decrements in areas such as the ACC and NAc (52). Thus, the absence of change in FC of the NAc with other parts of the reward circuit could suggest that downregulation of CB1 receptors mitigate the impact of acute cannabis intoxication on neural activity within fronto-subcortical circuits and associated behavioural outcomes, thus demonstrating the prime mechanisms underlying the development of tolerance in this circuit. However, to further explore the association between CB1 receptor availability and development of tolerance, future studies should assess CB1 receptor availability during acute intoxication.

In order to assess potential long-term effects of repeated cannabis exposure, analyses were performed comparing the placebo conditions between CUs and OUs on ROI-to-ROI FC within the reward circuit, metabolite concentration levels, and performance on the sustained attention task. Furthermore, as CUs had significantly higher baseline serum THC and THC-metabolite levels than OUs, these values were added as covariates. When controlling for serum concentration levels, no differences were seen between groups on any of the outcome variables. Absence of group differences are in line with previous PET studies which have found CB1R availability normalization in cannabis dependent users after as little as 2 days of monitored abstinence (51, 53), as well neuropsychological data suggesting reversible cognitive deficits, modulated more by recent exposure than by cumulative lifetime use (55). However, the literature on long-lasting effects of cannabis use is mixed, with studies also reporting long-term changes on brain structure (22, 23), neurometabolite concentration levels (56), and neurocognitive functioning (57). Importantly, results have been found to be region and domain specific, and influenced by factors such as frequency and age of onset of use, potentially explaining the variance in reported outcomes. In order to further assess chronic effects of repeated cannabis exposure on the reward system and associated behaviour, future studies should employ a larger sample size and take such factors into account.

Finally, MRS demonstrated cannabis induced changes in neurometabolites previously found to be altered in cannabis users (56), namely NAA and mI. NAA and mI, as well as glutamate, are markers of glial and neuronal activation (58, 59). Our data demonstrate that cannabis acutely increases these metabolites in the striatum in OUs. Treatment induced changes in NAA and mI also correlated with behavioural changes. Specifically, it was found that cannabis induced changes in mI in the striatum negatively correlated with cannabis induced changes in subjective high. Interestingly, elevations in mI and glutamate have been found in individuals with first-episode psychosis, and were found to correlate with subjective reports of grandiosity (60). Furthermore, in the present study it was found that cannabis induced changes in NAA positively correlated with cannabis induced changes in sustained attention performance. Similarly, altered levels of NAA and NAAG have been reported in patients with schizophrenia, albeit in the ACC, with NAA levels correlating with attention performance (61). Taken together, results suggest that cannabis increases glial and neuronal activation in the striatum, which is associated with changes in subjective state and behavioural performance. Importantly, these changes were not seen in the CUs, further demonstrating pharmacodynamic tolerance. However, cannabis decreased mI in the ACC in CUs, but not OUs. The finding of acute decrease in ACC mI is compatible with previous studies which found decreased mI in the ACC (19, 20), as well as throughout the brain (26, 27), and has been suggested to reflect cannabis-related immunosuppression (56).

In summary, our study provides previously unidentified evidence to suggest that a reduced responsiveness of the reward circuitry underlies a blunted pharmacodynamic response to an acute cannabis challenge in chronic users. Understanding the neuroadaptive basis of tolerance is important in the context of the therapeutic use of cannabis-based medications, as well as in the context of public health and safety of cannabis use when performing day to day operations.

## References

1. UN (2018) World Drug Report 2018. United Nations publication.

2. EMCDDA (2018) European Drug Report 2018: Trends and Developments. Publications Office of the European Union, Luxemborg.

3. Colizzi M & Bhattacharyya S (2018) Cannabis use and the development of tolerance: a systematic review of human evidence. Neurosci Biobehav Rev 93:1–25.

4. Iversen L (2003) Cannabis and the brain. Brain : a journal of neurology 126(Pt 6):1252–1270.

5. Diana M, Melis M, & Gessa G (1998) Increase in meso-prefrontal dopaminergic activity after stimulation of CB1 receptors by cannabinoids. Eur J Neurosci 10(9):2825–2830.

6. Fadda P, et al. (2006) Cannabinoid self-administration increases dopamine release in the nucleus accumbens. Neuroreport 17(15):1629–1632.

7. Ton JMNC, et al. (1988) The effects of Δ9-tetrahydrocannabinol on potassium-evoked release of dopamine in the rat caudate nucleus: an in vivo electrochemical and in vivo microdialysis study. Brain research 451(1-2):59–68.

8. Bossong MG, et al. (2015) Further human evidence for striatal dopamine release induced by administration ofΔ 9-tetrahydrocannabinol (THC): selectivity to limbic striatum. Psychopharmacology 232(15):2723–2729.

9. Pistis M, et al. (2002) Delta(9)-tetrahydrocannabinol decreases extracellular GABA and increases extracellular glutamate and dopamine levels in the rat prefrontal cortex: an in vivo microdialysis study. Brain Res 948(1-2):155–158.

10. Mason NL, et al. (2018) Cannabis induced increase in striatal glutamate associated with loss of functional corticostriatal connectivity. European Neuropsychopharmacology 29(2):247–256.

11. Colizzi M, et al. (2019) Delta-9-tetrahydrocannabinol increases striatal glutamate levels in healthy individuals: implications for psychosis. Molecular psychiatry.

12. Scofield MD, et al. (2016) The Nucleus Accumbens: Mechanisms of Addiction across Drug Classes Reflect the Importance of Glutamate Homeostasis. Pharmacological reviews 68(3):816–871.

13. Peter W. Kalivas, Ph.D., and & Nora D. Volkow, M.D. (2005) The Neural Basis of Addiction: A Pathology of Motivation and Choice. American Journal of Psychiatry 162(8):1403–1413.

14. van de Giessen E, et al. (2017) Deficits in striatal dopamine release in cannabis dependence. Molecular psychiatry 22(1):68.

15. Bloomfield MAP, et al. (2014) Dopaminergic Function in Cannabis Users and Its Relationship to Cannabis-Induced Psychotic Symptoms. Biological psychiatry 75(6):470–478.

16. Leroy C, et al. (2012) Striatal and extrastriatal dopamine transporter in cannabis and tobacco addiction: a high-resolution PET study. Addiction biology 17(6):981–990.

17. Muetzel RL, et al. (2013) In vivo 1H magnetic resonance spectroscopy in young-adult daily marijuana users. NeuroImage: Clinical 2:581–589.

18. Chang L, Cloak C, Yakupov R, & Ernst T (2006) Combined and independent effects of chronic marijuana use and HIV on brain metabolites. Journal of neuroimmune pharmacology : the official journal of the Society on NeuroImmune Pharmacology 1(1):65–76.

19. Prescot AP, Locatelli AE, Renshaw PF, & Yurgelun-Todd DA (2011) Neurochemical alterations in adolescent chronic marijuana smokers: a proton MRS study. NeuroImage 57(1):69–75.

20. Prescot AP, Renshaw PF, & Yurgelun-Todd DA (2013) gamma-Amino butyric acid and glutamate abnormalities in adolescent chronic marijuana smokers. Drug and alcohol dependence 129(3):232–239.

21. Hill SY, Sharma V, & Jones BL (2016) Lifetime use of cannabis from longitudinal assessments, cannabinoid receptor (CNR1) variation, and reduced volume of the right anterior cingulate. Psychiatry Research: Neuroimaging 255:24–34.

22. Filbey FM, et al. (2014) Long-term effects of marijuana use on the brain. Proceedings of the National Academy of Sciences 111(47):16913–16918.

23. Battistella G, et al. (2014) Long-term effects of cannabis on brain structure. Neuropsychopharmacology 39(9):2041–2048.

24. Hermann D, et al. (2007) Dorsolateral Prefrontal Cortex N-Acetylaspartate/Total Creatine (NAA/tCr) Loss in Male Recreational Cannabis Users. Biological psychiatry 61(11):1281–1289.

25. Cowan RL, Joers JM, & Dietrich MS (2009) N-acetylaspartate (NAA) correlates inversely with cannabis use in a frontal language processing region of neocortex in MDMA (Ecstasy) polydrug users: a 3 T magnetic resonance spectroscopy study. Pharmacology Biochemistry and Behavior 92(1):105–110.

26. Mashhoon Y, Jensen JE, Sneider JT, Yurgelun-Todd DA, & Silveri MM (2013) Lower left thalamic myo-inositol levels associated with greater cognitive impulsivity in marijuana-dependent young men: preliminary spectroscopic evidence at 4T. Journal of addiction research & therapy.

27. Silveri MM, Jensen JE, Rosso IM, Sneider JT, & Yurgelun-Todd DA (2011) Preliminary evidence for white matter metabolite differences in marijuana-dependent young men using 2D J-resolved magnetic resonance spectroscopic imaging at 4 Tesla. Psychiatry research 191(3):201–211.

28. Sung YH, et al. (2013) Decreased frontal N-acetylaspartate levels in adolescents concurrently using both methamphetamine and marijuana. Behav Brain Res 246:154–161.

29. Ramaekers J, et al. (2013) Methylphenidate reduces functional connectivity of nucleus accumbens in brain reward circuit. Psychopharmacology (Berl) 229(2):219–226.

30. Ramaekers JG, et al. (2016) Cannabis and cocaine decrease cognitive impulse control and functional corticostriatal connectivity in drug users with low activity DBH genotypes. Brain Imaging and Behavior 10(4):1254–1263.

31. Ramaekers JG, Kauert G, Theunissen E, Toennes SW, & Moeller M (2009) Neurocognitive performance during acute THC intoxication in heavy and occasional cannabis users. Journal of psychopharmacology 23(3):266–277.

32. Hartley S, et al. (2019) Effect of Smoked Cannabis on Vigilance and Accident Risk Using Simulated Driving in Occasional and Chronic Users and the Pharmacokinetic–Pharmacodynamic Relationship. Clinical chemistry:clinchem. 2018.299727.

33. Theunissen EL, et al. (2012) Neurophysiological functioning of occasional and heavy cannabis users during THC intoxication. Psychopharmacology (Berl) 220(2):341–350.

34. Ramaekers JG, et al. (2016) Cannabis and tolerance: acute drug impairment as a function of cannabis use history. Scientific Reports 6.

35. Frahm J, et al. (1989) Localized high-resolution proton NMR spectroscopy using stimulated echoes: initial applications to human brain in vivo. Magnetic resonance in medicine 9(1):79–93.

36. Chao-Gan Y & Yu-Feng Z (2010) DPARSF: A MATLAB Toolbox for “Pipeline” Data Analysis of Resting-State fMRI. Frontiers in systems neuroscience 4:13.

37. Dinges DF & Powell JW (1985) Microcomputer analyses of performance on a portable, simple visual RT task during sustained operations. Behavior Research Methods, Instruments, & Computers 17(6):652–655.

38. Toennes SW, Ramaekers JG, Theunissen EL, Moeller MR, & Kauert GF (2008) Comparison of cannabinoid pharmacokinetic properties in occasional and heavy users smoking a marijuana or placebo joint. Journal of analytical toxicology 32(7):470–477.

39. Cummings JL (1993) Frontal-subcortical circuits and human behavior. Archives of neurology 50(8):873–880.

40. Alexander GE, DeLong MR, & Strick PL (1986) Parallel organization of functionally segregated circuits linking basal ganglia and cortex. Annual review of neuroscience 9(1):357–381.

41. Bonelli RM & Cummings JL (2007) Frontal-subcortical circuitry and behavior. Dialogues in clinical neuroscience 9(2):141.

42. Perreault ML, Hasbi A, O’Dowd BF, & George SR (2011) The dopamine D1–D2 receptor heteromer in striatal medium spiny neurons: evidence for a third distinct neuronal pathway in basal ganglia. Frontiers in neuroanatomy 5:31.

43. Pierce RC & Kumaresan V (2006) The mesolimbic dopamine system: the final common pathway for the reinforcing effect of drugs of abuse? Neurosci Biobehav Rev 30(2):215–238.

44. Volkow ND, Wang GJ, Fowler JS, Tomasi D, & Telang F (2011) Addiction: beyond dopamine reward circuitry. Proc Natl Acad Sci U S A 108(37):15037–15042.

45. Lupica CR, Riegel AC, & Hoffman AF (2004) Marijuana and cannabinoid regulation of brain reward circuits. British journal of pharmacology 143(2):227–234.

46. Breivogel CS, et al. (1999) Chronic delta9-tetrahydrocannabinol treatment produces a time-dependent loss of cannabinoid receptors and cannabinoid receptor-activated G proteins in rat brain. Journal of neurochemistry 73(6):2447–2459.

47. Sim-Selley LJ (2003) Regulation of cannabinoid CB1 receptors in the central nervous system by chronic cannabinoids. Critical reviews in neurobiology 15(2):91–119.

48. McKinney DL, et al. (2008) Dose-related differences in the regional pattern of cannabinoid receptor adaptation and in vivo tolerance development to delta9-tetrahydrocannabinol. The Journal of pharmacology and experimental therapeutics 324(2):664–673.

49. Sim LJ, Hampson RE, Deadwyler SA, & Childers SR (1996) Effects of chronic treatment with delta9-tetrahydrocannabinol on cannabinoid-stimulated [35S]GTPgammaS autoradiography in rat brain. The Journal of neuroscience : the official journal of the Society for Neuroscience 16(24):8057–8066.

50. Romero J, et al. (1997) Atypical location of cannabinoid receptors in white matter areas during rat brain development. Synapse 26(3):317–323.

51. Hirvonen J, et al. (2012) Reversible and regionally selective downregulation of brain cannabinoid CB1 receptors in chronic daily cannabis smokers. Molecular psychiatry 17(6):642–649.

52. Ceccarini J, et al. (2015) [18F]MK-9470 PET measurement of cannabinoid CB1 receptor availability in chronic cannabis users. Addict Biol 20(2):357–367.

53. D’Souza DC, et al. (2016) Rapid Changes in CB1 Receptor Availability in Cannabis Dependent Males after Abstinence from Cannabis. Biological psychiatry. Cognitive neuroscience and neuroimaging 1(1):60–67.

54. Gonzalez S, Cebeira M, & Fernandez-Ruiz J (2005) Cannabinoid tolerance and dependence: a review of studies in laboratory animals. Pharmacology, biochemistry, and behavior 81(2):300–318.

55. Pope HG, Jr, Gruber AJ, Hudson JI, Huestis MA, & Yurgelun-Todd D (2001) Neuropsychological Performance in Long-term Cannabis Users. JAMA Psychiatry 58(10):909–915.

56. Sneider JT, Mashhoon Y, & Silveri MM (A Review of Magnetic Resonance Spectroscopy Studies in Marijuana using Adolescents and Adults. Journal of addiction research & therapy Suppl 4.

57. Crean RD, Crane NA, & Mason BJ (2011) An evidence based review of acute and long-term effects of cannabis use on executive cognitive functions. Journal of addiction medicine 5(1):1–8.

58. Harte MK, Bachus SB, & Reynolds GP (2005) Increased N-acetylaspartate in rat striatum following long-term administration of haloperidol. Schizophrenia research 75(2-3):303–308.

59. Malhi GS, Valenzuela M, Wen W, & Sachdev P (2002) Magnetic resonance spectroscopy and its applications in psychiatry. The Australian and New Zealand journal of psychiatry 36(1):31–43.

60. Plitman E, et al. (2016) Elevated Myo-Inositol, Choline, and Glutamate Levels in the Associative Striatum of Antipsychotic-Naive Patients With First-Episode Psychosis: A Proton Magnetic Resonance Spectroscopy Study With Implications for Glial Dysfunction. Schizophrenia bulletin 42(2):415–424.

61. Jessen F, et al. (2013) N-acetylaspartylglutamate (NAAG) and N-acetylaspartate (NAA) in patients with schizophrenia. Schizophrenia bulletin 39(1):197–205.

