## Supplemental Information for "Reduced responsiveness of the reward system underlies tolerance to cannabis impairment in chronic users"

Natasha L Mason

**This PDF file includes:**

Supplementary text

8 Tables

2 Figures

References for SI reference citations

### Supplementary Information Text

### SI Methods

**Participants.** Participants were recruited through advertisements around Maastricht University. Inclusion criteria were: age, 18-40 years; occasional cannabis use for the OU group, ranging between 1 time a month and 3 times a week for the past year OR chronic cannabis use for the CU group, using at least 4 times a week for the past year; normal weight, body mass index between 18 and 28 kg/m2; free from psychotropic medication; good physical health, including absence of major medical, endocrine, and neurological conditions; and written informed consent. Exclusion criteria were: history of drug abuse (other than the use of cannabis) or addiction, which were determined by medical questionnaires and examination; pregnancy or lactation; health issues including hypertension (diastolic >90 and systolic >140), cardiac dysfunction, and liver dysfunction; current or history of psychiatric disorders; previous experience of serious side effects to cannabis; and MRI contraindications. Before inclusion, subjects were screened and examined by a study physician, who checked for general health, conducted a resting ECG, and took blood and urine samples in which hematology, clinical chemistry, urine, and virology analyses were conducted. Participant demographic data can be found in Table S1.

A permit for obtaining, storing, and administering cannabis was obtained from the Dutch Drug Enforcement Administration. Participants were financially compensated for their participation in the study.

**Administration.** Both treatments were administered through a Volcano vaporizer (Storz & Bickel Volcano ®), with participants inhaling equal amounts of each while lying in the MRI scanner. The treatments were vaporized at 225°C and the vapor was stored in a polythene bag equipped with a mouth piece. Participants were instructed to place the mouth piece to their lips and inhale deeply, holding their breath for 10 seconds, and then exhaling. Participants repeated this procedure until balloon was empty. Participants were instructed to inhale the entire volume of the balloon within 5 minutes, according to a standardized procedure.

**Procedures.** Participants were familiarized with the test day procedures on a separate training day prior to the treatment conditions. Participants in the OU group were instructed to refrain from drug use, including cannabis, (≥7 days) and alcohol use (≥24 hours) prior to their testing day; whereas participants in the CU group were given the same instructions, however were allowed to use cannabis up until 24 hours prior to their testing day. On arrival on a test day, absence of drug and alcohol were assessed via a urine drug screen and a breath alcohol screen. An additional pregnancy test was given if participants were female. If all tests were found to be negative (except for cannabis in the CU group), participants were allowed to proceed, and a venal catheter was placed.

**Psychomotor vigilance task.** This study used a shortened, 5-minute version of the task on three consecutive time points after treatment administration. The outcome variables of the PVT are: mean RT, the mean RT for all trials; optimum response domain, the fastest 10% of response times for all trials (i.e. average of the fastest 10% RT); number of lapses, the number of response times greater than or equal to 500 ms for all trials (i.e. lapse); and lapse domain, the slowest 10%

of reciprocal response times for all trials (i.e. average of the slowest 10% 1/RT) (1). For this study, only number of lapses was considered.

**MRS Data Acquisition and Quality.** Single-voxel proton magnetic resonance spectroscopy (MRS) measurements were performed on a MAGNETOM 7T MR scanner (Siemens Healthineers, Erlangen, Germany) with a whole-body gradient set (SC72; maximum amplitude, 70 mT/m; maximum slew rate, 200 T/m/s) and using an single-channel transmit/32-channel receive head coil (Nova Medical, Wilmington, MA, USA). Spectroscopic voxels of interest were placed by a trained operator at the Anterior Cingulate Cortex (ACC) (voxel size = 25 x 20 x 17 $mm^3$) and the right striatum (voxel size = 20 x 20 x 20 $mm^3$). Spectra were acquired with stimulated echo acquisition mode (STEAM) (2) sequence using the following parameters: TE = 6.0 ms, TM = 10.0 ms, TR = 5.0 s, NA = 64, flip angle = 90°, RF bandwidth = 4.69 kHz, RF centred at 2.4 ppm, receive bandwidth = 4.0 kHz, vector size = 2048, 16-step phase cycling, acquisition time = 5:20 min. Water suppression was achieved by variable power RF pulses with optimised relaxation delays (VAPOR) (3). In addition, a complete phase cycle of measurements was acquired without the water suppression RF pulses to record a water peak reference for eddy current correction (4) and absolute metabolite concentration calibration (5, 6). Before the spectroscopy measurements, a 3D-GRE dual-echo field-map ($TE_1$ = 1.00 ms, $TE_2$ = 2.98 ms, TR = 20.0 ms, flip angle = 8°, voxel size = 3 mm isotropic, matrix size = 84 × 84 × 56, bandwidth = 1450 Hz/pixel, acquisition time = 2:24 min) was acquired and used calculate the shim currents required to homogenise the static magnetic field in the spectroscopic voxels of interest.

The spectra were analysed with LCModel version 6.3-1H using a GAMMA (7) simulated basis set which includes Alanine (Ala), Ascorbic Acid (Asc), Aspartate (Asp), Creatine (Cr), γ-Aminobutyric Acid (GABA), Glucose (Glc), Glutamate (Glu), Glutamine (Gln), Glycerophosphocholine (GPC), Glutathione (GSH), Glycine (Glyc), Lactate (Lac), Myo-Inositol (mI), N-Acetyl Aspartate (NAA), N-Acetyl Aspartyl Glutamate (NAAG), Phosphocreatine (PCr), Phosphorylcholine (PCh), Phosphorylethanolamine (PE) , Scyllo-Inositol (Scyllo), and Taurine (Tau) (8). The metabolite basis set also includes an in vivo Macromolecules (MMol) spectrum which was collected using a metabolites suppressed double inversion recovery (DIR) STEAM with the same parameters as above and $TI_1$ = 2.09 s and $TI_2$ =0.52 s (9).

Anatomical ($T_1$-weighted) images were acquired using magnetisation-prepared 2 rapid acquisition gradient-echo (MP2RAGE) (10) sequence (TR = 4.5 s, TE = 2.39 ms, $TI_1$ = 0.90 s, $TI_2$ = 2.75 s, flip $angle_1$ = 5°, flip $angle_2$ = 3°, voxel size = 0.9 mm isotropic, matrix size = 256 × 256 × 192, phase partial Fourier = 6/8, GRAPPA factor = 3 with 24 reference lines, bandwidth = 250 Hz/pixel, acquisition time = 6:00 min). Tissue probability maps for grey matter (GM), white matter (WM) and cerebrospinal fluid (CSF) were generated from the $T_1$-weighted anatomical images using FSL-FAST (11). To ensure data quality and reliable metabolite estimation, only absolute metabolite values with a Cramer–Rao lower bound below 20% and a signal-to-noise ratio greater than 10 were considered (Provencher, 2001). An example spectrum can be found in Figure S2, and mean SNR and %CRLB values in Table S6.

**fMRI data preprocessing and functional connectivity.** Resting state image preprocessing were conducted using SPM8 (Statistical Parametric Mapping, Wellcome Trust Centre for

Neuroimaging, Institute of Neurology, University College London) Preprocessing steps included: motion correction (registered to the first image with second degree B-spline interpolation), coregistration (linking of functional to anatomical scans), and spatial normalization (the mean EPI image of each session was matched to SPM8's EPI template in Montreal Neurological Institute [MNI] space) where after the parameters were applied to all images of that session. During normalization voxel size was 1.5 x 1.5 x 1.5 mm. Finally, the data were smoothed at 3mm Gaussian kernel.

Linear trends of time courses were removed followed by low band-pass filtered (0.01-0.08 HZ) of the preprocessed data to remove 'noise' attributable to physiological parameters. Nuisance covariates (motion parameters, white matter signal, CSF signal) were also removed. In order to indirectly assess dopamine neurotransmission, two spheres (4 mm radius) were created that were located (in MNI space) in the left (-9, 9, -9) and right (9, 9, -9) NAc. Average time courses were obtained for each sphere separately and correlational analysis was performed voxel wise to generate functional connectivity maps for each sphere. The correlation coefficient map was converted into z maps by Fisher's r-to-z transform to improve normality (12, 13). This is in accordance with previous studies investigating drug induced changes in functional connectivity (14, 15).

Furthermore, as we were interested in FC *within* the reward circuit, ROI-to-ROI FC was computed according to the same aforementioned procedure, between areas including: NAc (-9, 9, -9, radius 4mm), MDN (-9, -19, -6, 4 mm), VP (-20, -4, -2, 4 mm) and the MC (-4, -18, 44, 4mm). NAc, MDN, and VP seed locations were in agreement with structural and functional subdivisions of these brain regions that were validated in previous work (16-18). The MC was based off of the first FC analysis. Analysis was performed between the

NAc and MDN, NAc and VPN, MDN and VPN, MDN and MC, and MCand NAc. Treatment change was averaged across the 2 time points [Average(THC time point 1- placebo time point 1); (THC time point 2 – placebo time point 2)].

**Statistical analysis of FC data.** Functional connectivity data (i.e. correlation coefficient maps for each individual in each treatment condition at each time point) were analyzed in a GLM model in SPM 12. In the first GLM, data entered a full factorial model with treatment (THC and placebo) and time point (2 levels) as within-subject factors and group (occasional and chronic user group) as a between-subject factor. From this model main effects of treatment were identified for the occasional user group, but not the chronic user group. Maps were corrected for multiple comparisons at the cluster level using the family wise error correction (FWE).

**Correlational analyses.** Correlation analyses were conducted to further investigate the relationship between THC induced changes in brain and behavior. In order to reduce the number of comparisons, only variables significantly affected by THC were assessed (table S7).For the psychomotor vigilance task, only number of lapses was used as a variable of sustained attention. For the voxel wise correlation analysis (Table S8) between NAc FC and behavioral outcomes, individual treatment maps (placebo > THC) were entered into one-sample *t*-tests in SPM, with the average change scores of subjective high, and number of attention lapses. Maps were corrected for multiple comparisons at the cluster level using the family wise error correction (FWE). Average mean voxel activation of SPM identified clusters were put into SPSS and Pearson's correlations were performed to get correlation strengths.

### References


1. Loh S, Lamond N, Dorrian J, Roach G, & Dawson D (2004) The validity of psychomotor vigilance tasks of less than 10-minute duration. *Behavior research methods, instruments, & computers : a journal of the Psychonomic Society, Inc* 36(2):339-346.
2. Frahm J*, et al.* (1989) Localized high-resolution proton NMR spectroscopy using stimulated echoes: initial applications to human brain in vivo. *Magnetic resonance in medicine* 9(1):79-93.
3. Tkac I, Starcuk Z, Choi IY, & Gruetter R (1999) In vivo 1H NMR spectroscopy of rat brain at 1 ms echo time. *Magnetic resonance in medicine* 41(4):649-656.
4. Klose U (1990) In vivo proton spectroscopy in presence of eddy currents. *Magnetic resonance in medicine* 14(1):26-30.
5. Barker PB*, et al.* (1993) Quantitation of proton NMR spectra of the human brain using tissue water as an internal concentration reference. *NMR in biomedicine* 6(1):89-94.
6. Soher BJ, Hurd RE, Sailasuta N, & Barker PB (1996) Quantitation of automated single-voxel proton MRS using cerebral water as an internal reference. *Magnetic resonance in medicine* 36(3):335-339.
7. Smith SA, Levante TO, Meier BH, & Ernst RR (1994) Computer Simulations in Magnetic Resonance. An Object-Oriented Programming Approach. *Journal of Magnetic Resonance* 106(1):75-105.
8. Govindaraju V, Young K, & Maudsley AA (2000) Proton NMR chemical shifts and coupling constants for brain metabolites. *NMR in biomedicine* 13(3):129-153.
9. Penner J & Bartha R (2014) Semi-LASER 1 H MR spectroscopy at 7 Tesla in human brain: Metabolite quantification incorporating subject-specific macromolecule removal. *Magnetic resonance in medicine*.
10. Marques JP*, et al.* (2010) MP2RAGE, a self bias-field corrected sequence for improved segmentation and T1-mapping at high field. *NeuroImage* 49(2):1271-1281.
11. Zhang Y, Brady M, & Smith S (2001) Segmentation of brain MR images through a hidden Markov random field model and the expectation-maximization algorithm. *IEEE transactions on medical imaging* 20(1):45-57.
12. Chao-Gan Y & Yu-Feng Z (2010) DPARSF: A MATLAB Toolbox for "Pipeline" Data Analysis of Resting-State fMRI. *Frontiers in systems neuroscience* 4:13.
13. Rosner B (2006) *Fundamentals of Biostatistics* (Thomson-Brooks/Cole, Belmont, CA) 6 Ed.
14. Ramaekers J*, et al.* (2013) Methylphenidate reduces functional connectivity of nucleus accumbens in brain reward circuit. *Psychopharmacology (Berl)* 229(2):219-226.
15. Ramaekers JG*, et al.* (2016) Cannabis and cocaine decrease cognitive impulse control and functional corticostriatal connectivity in drug users with low activity DBH genotypes. *Brain Imaging and Behavior* 10(4):1254-1263.
16. Filbey FM, Aslan S, Lu H, & Peng S-L (2018) Residual effects of THC via novel measures of brain perfusion and metabolism in a large group of chronic cannabis users. *Neuropsychopharmacology* 43(4):700.
17. Di Martino A*, et al.* (2008) Functional connectivity of human striatum: a resting state FMRI study. *Cerebral cortex (New York, N.Y. : 1991)* 18(12):2735-2747.
18. Kelly C*, et al.* (2009) L-dopa modulates functional connectivity in striatal cognitive and motor networks: a double-blind placebo-controlled study. *The Journal of neuroscience : the official journal of the Society for Neuroscience* 29(22):7364-7378.

Table S1. Mean subject characteristics (SD) and history of drug use for occasional and chronic cannabis users that completed the study (N=24).

| Variable | Occasional Users | Chronic Users | Value | df | *P* value |
|---|---|---|---|---|---|
| Gender (male/female), n, total | 5/7, 12 | 9/3, 12 | $\chi2$ = 2.74‡ | 1 | 0.10 |
| Age, years | 22.5 (2.54) | 21.83 (2.25) | *t*=.681† | 22 | 0.50 |
| History of cannabis use, years | 5.50 (2.71) | 5.33 (1.78) | *t*=.178† | 22 | 0.86 |
| Frequency of cannabis use, per week | 1.21 (0.80) | 6.63 (1.40) | *t*=-11.62† | 22 | 0.00* |
| Alcohol consumption, glasses per week | 7.29 (7.05) | 3.17 (2.32) | *t*=1.93† | 22 | 0.07 |
| Caffeine consumption (per week) | 8.25 (7.71) | 8.88 (6.05) | *t*=-0.22† | 22 | 0.83 |
| Nicotine consumption, per week | 16.33 (22.07) | 19.29 (28.23) | *t*=-1.25† | 22 | 0.22 |
| Occasional use of other drugs, n | 9 | 9 | $\chi2$ = 0.00‡ | 1 | 1.00 |

*Significant *P* values

†Independent *t* test

‡$\chi2$ test for frequency data

Table S2. Time course for mean (S.E.) concentrations of THC and its metabolites in serum (ng/ml) following smoking THC, as assessed by gas chromatography-mass spectrometry (GC-MS).

| Time Relative to Smoking | Serum (GC-MS) | | | | | |
|---|---|---|---|---|---|---|
| | Occasional Users | | | Chronic Users | | |
| | THC | 11-OH-THC | THC-COOH | THC | 11-OH-THC | THC-COOH |
| 0 | 0.00 (.00)* | 0.02 (.02) | 1.48 (0.71) | 3.48 (.89) | 1.55 (.38) | 47.44 (14.33) |
| 6 | 7.80 (1.69) | 1.70 (0.43) | 9.50 (2.23) | 15.86 (3.48) | 3.84 (1.13) | 48.81 (14.19) |
| 28 | 2.86 (0.55) | 1.03 (0.22) | 8.37 (1.82) | 6.66 (1.55) | 2.10 (.62) | 45.53 (15.43) |
| 50 | 2.07 (0.34) | 0.86 (0.20) | 7.55 (1.55) | 7.10 (1.44) | 2.07 (.46) | 45.72 (12.23) |
| 67 | 2.22 (0.52) | 1.00 (0.23) | 8.60 (1.94) | 5.67 (1.48) | 1.67 (.47) | 36.90 (13.27) |

*one participant exhibited a THC concentration of 1.1 ng/ml, which was not regarded as indicative of recent use due to a low THC-OH concentration (0.2 ng/ml).

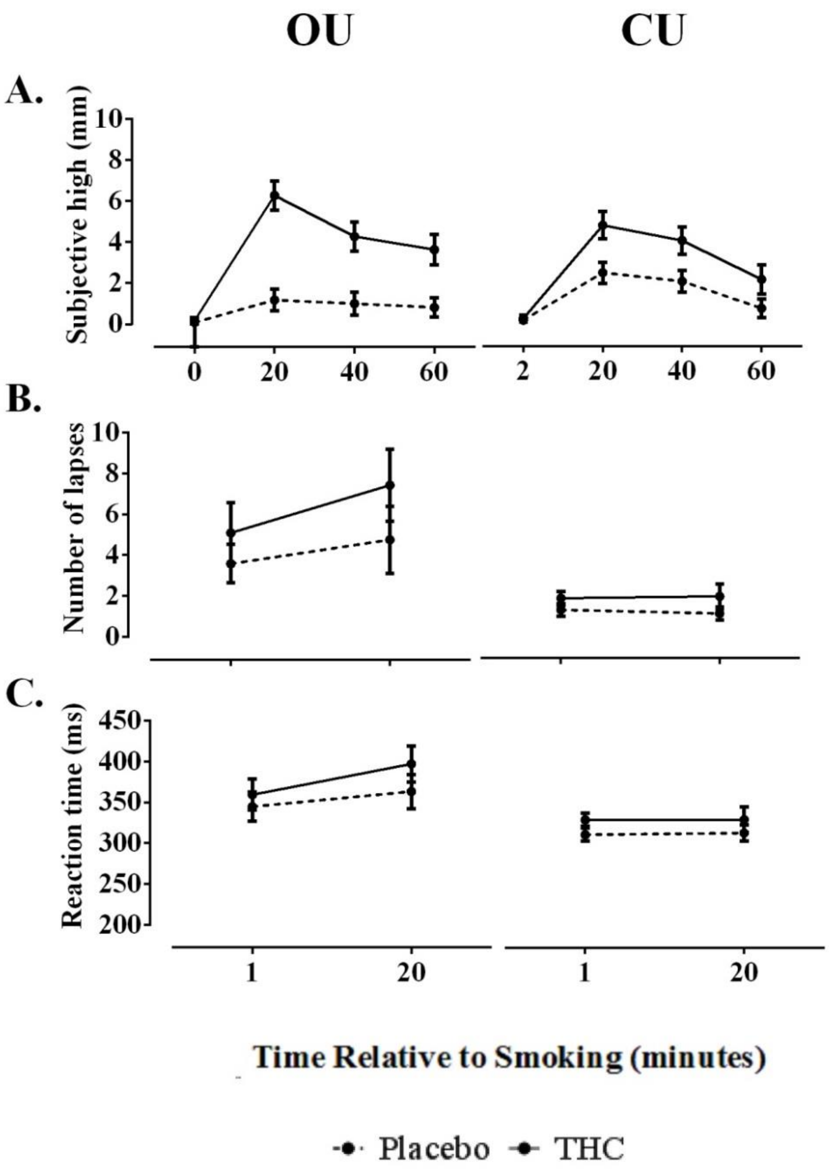


Figure S1. Subjective and cognitive effects. A. Occasional and chronic users mean (SE) subjective high for both treatments (THC vs placebo) as a function of time relative to smoking (0, 20, 40, and 60 minutes). B. Occasional and chronic users mean (SE) reaction time, and C. number of lapses (RT>500 ms) for both treatments (THC vs placebo) as a function of time relative to smoking (1, 20, and 40 minutes).

Table S3. Mean (SE) metabolite concentrations for OU and CU's.

| | | Glu/tCr | | GABA/tCr | | NAA + NAAG/tCr | | mI/tCr | | GPC + PCh/tCr | |
|---|---|---|---|---|---|---|---|---|---|---|---|
| | | Occasional Users | | | | | | | | | |
| | Time | THC | Placebo | THC | Placebo | THC | Placebo | THC | Placebo | THC | Placebo |
| Anterior Cingulate Cortex | 1 | 1.04 (.05) | 1.05 (.03) | .18 (.01) | .17 (.04) | 1.15 (.06) | 1.20 (.05) | .87 (.02) | .82 (.03) | 0.18 (.01) | .19 (.01) |
| | 2 | 1.04 (.04) | 1.05 (.03) | .15 (.01) | .14 (.01) | 1.17 (.05) | 1.17 (.04) | .86 (.02) | .81 (.03) | 0.17 (.01) | .19 (.01) |
| Striatum | 1 | 1.03 (.04) | .99 (.05) | .20 (.01) | .19 (.02) | 1.45 (.05) | 1.36 (.06) | .58 (.03) | .49 (.03) | .15 (.01) | .16 (.01) |
| | 2 | 1.14 (.05) | .93 (.02) | .30 (.04) | .19 (.03) | 1.58 (.06) | 1.34 (.05) | .61 (.04) | .52 (.03) | .16 (.01) | .15 (.01) |
| Chronic Users | | | | | | | | | | | |
| Anterior Cingulate Cortex | 1 | 1.06 (.02) | 1.05 (.06) | .15 (.01) | .17 (.03) | 1.18 (.03) | 1.19 (.06) | .76 (.05) | .90 (.05) | .18 (.01) | .19 (.01) |
| | 2 | 1.08 (.03) | 1.05 (.05) | .15 (.01) | .15 (.01) | 1.25 (.03) | 1.24 (.04) | .79 (.04) | .89 (.85) | .18 (.01) | .18 (.01) |
| Striatum | 1 | 1.02 (.04) | 1.08 (.07) | .24 (.03) | .26 (.02) | 1.36 (.08) | 1.34 (.05) | .56 (.05) | .57 (.05) | .14 (.01) | .16 (.01) |
| | 2 | .97 (.03) | 1.07 (.06) | .20 (.02) | .24 (.04) | 1.44 (.05) | 1.35 (.05) | .50 (.03) | .53 (.03) | .16 (.01) | .15 (.01) |

Table S4. Significant decrements in functional connectivity of left nucleus accumbens, relative to placebo in the occasional group.

| Cluster | R or L | BA | k | x | y | z | P value $t_{3.20}$ FWE cluster corrected |
|---|---|---|---|---|---|---|---|
| Precuneus | L | 5 | 71 | -6 | -50 | 56 | .000 |
| Middle Frontal Gyrus | R | 46 | 22 | 34 | 38 | 30 | .016 |
| Middle Frontal Gyrus | L | 6 | 19 | -38 | 6 | 62 | .043 |
| Superior Frontal Gyrus | L | 6 | 48 | -24 | 6 | 64 | .000 |
| Precentral Gyrus | L | 6 | 39 | -28 | -26 | 68 | .000 |
| Precentral Gyrus | L | 6 | 39 | -48 | -4 | 42 | .000 |
| Precentral Gyrus | R | 6 | 24 | 62 | 8 | 18 | .000 |
| Precentral Gyrus | R | 6 | 24 | 54 | -2 | 46 | .008 |
| Supplementary Motor Area | L | 6 | 24 | -6 | 8 | 48 | .008 |
| Supplementary Motor Area | R | 6 | 24 | 6 | 12 | 66 | .008 |
| Supplementary Motor Area | R | 6 | 44 | 8 | -6 | 68 | .000 |
| Rolandic Operculum | R | 6 | 44 | 62 | 8 | 10 | .000 |
| Rolandic Operculum | L | 48 | 108 | -58 | 4 | 4 | .000 |
| | L | 48 | 115 | -36 | 18 | 20 | .000 |
| | L | 48 | 20 | -34 | 8 | 0 | .000 |
| | L | 48 | 21 | -30 | -30 | 12 | .022 |
| | R | 48 | 52 | 44 | 12 | 18 | .000 |
| Midcingulate Area | R | 23 | 96 | 6 | -24 | 46 | .000 |
| Midcingulate Area | L | 23 | 38 | -4 | -18 | 44 | .000 |

| | | | | | | | |
|---|---|---|---|---|---|---|---|
| Postcentral Gyrus | R | 43 | 57 | 60 | 2 | 24 | .000 |
| Postcentral Gyrus | L | 3 | 26 | -48 | -20 | 58 | .000 |
| Inferior Parietal Lobule | L | 2 | 28 | -48 | -40 | 58 | .002 |
| Inferior Parietal Lobule | L | 40 | 140 | -58 | -46 | 42 | .000 |
| Supramarginal Gyrus | L | 40 | 71 | -60 | -36 | 32 | .000 |
| Calcarine Sulcus | R | 17 | 36 | 18 | -62 | 6 | .000 |
| Middle Occipital Gyrus | L | 19 | 19 | -38 | -86 | 24 | .043 |
| | L | 19 | 36 | -40 | -88 | -14 | .000 |

Statistical threshold: $P < 0.001$ (uncorrected). MNI coordinates of peak voxels for each cluster are given.

Table S5. Testing day schedule

| Time after treatment (minutes) | Procedure |
|---|---|
| Baseline | |
| 0 | Urine sample (drug screen; pregnancy test) |
| 0 | Alcohol breath test |
| 0 | Vital signs |
| 0 | Questionnaires |
| 0 | Blood sample (S1) |
| In Scanner | |
| 0 | Anatomical scan |
| 0 | Administration (300 μg/kg) |
| 1 | MRS (ACC); PVT |
| 6 | Blood sample (S2) |
| 8 | MRS (striatum) |
| 15 | Resting state scan |
| 20 | Questionnaires |
| 22 | MRS (ACC); PVT |
| 28 | Blood sample (S3) |

| | |
|---|---|
| 30 | MRS (striatum) |
| 36 | Resting state scan |
| 42 | Questionnaires |
| 50 | Blood sample (S4) |
| Out of scanner | |
| 68 | Questionnaires |
| 72 | Blood sample (S5) |
| 75 | Sober up |

Figure S2. Example LC-Model fitted 1H-MRS data recorded from one participant. The black line spectra corresponds to the phased 1H-MRS data with the LC-Model fits overlaid (red). The residual spectra (raw data minus the LC-Model fit) are displayed below the spectrum.

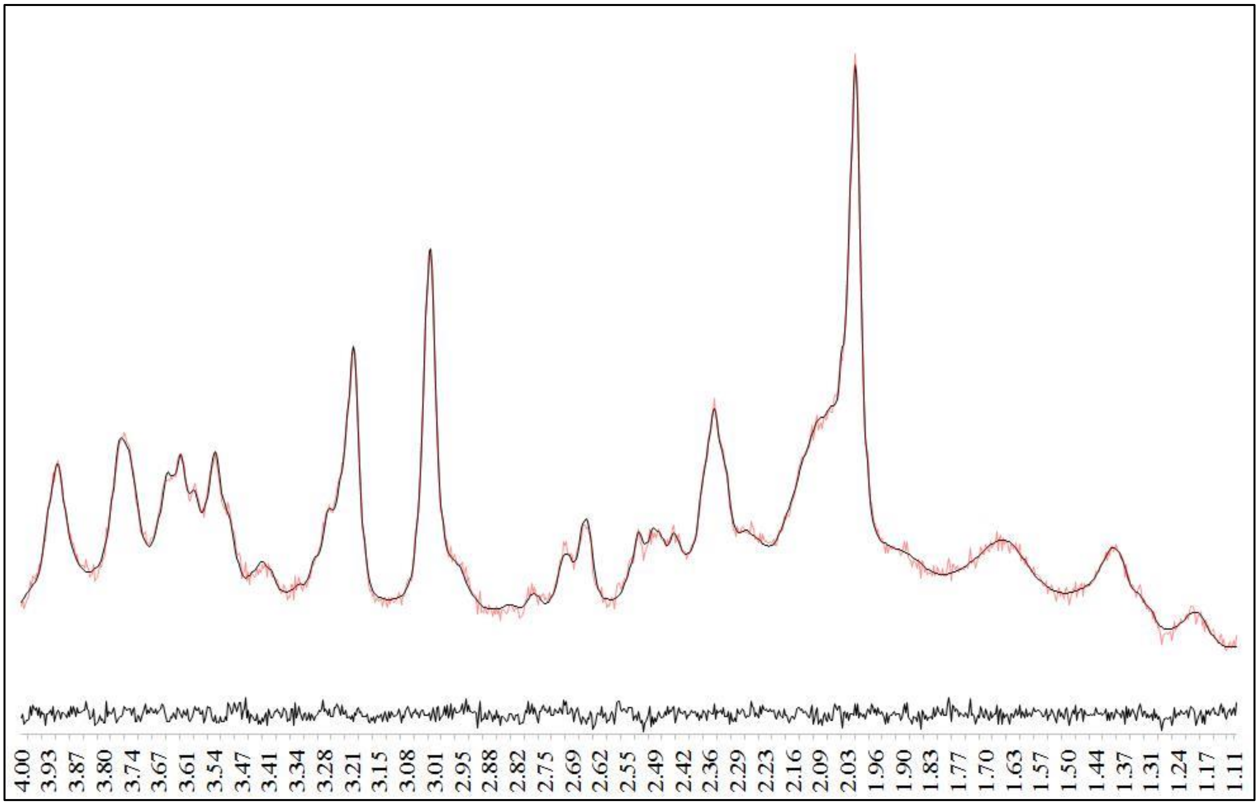

Table S6. Mean (min-max) ratios for signal to noise (SNR) and %CRLB per group, per condition, and per timepoint. Only data points with an SNR > 10 and a CRLB < 20% were included in the final analysis.

| | SNR | | | %CRLB | | | | | | | | | |
|---|---|---|---|---|---|---|---|---|---|---|---|---|---|
| | | | | Glu | | GABA | | mI | | tNAA | | tCho | |
| | Time | THC | Placebo | THC | Placebo | THC | Placebo | THC | Placebo | THC | Placebo | THC | Placebo |
| Occasional Users | | | | | | | | | | | | | |
| Anterior Cingulate Cortex | 1 | 32.1 (17-52) | 31.5 (13-47) | 2.6 (2-4) | 2.9 (2-5) | 10.6 (9-18) | 13 (8-19) | 3.2 (2-6) | 41 (2-7) | 2.2 (2-3) | 2.3 (2-3) | 3.8 (2-8) | 4 (2-14) |
| | 2 | 31.1 (14-49) | 32.3 (15-48) | 2.9 (2-6) | 2.7 (2-4) | 12 (7-17) | 14.9 (7-19) | 3.4 (2-8) | 3.9 (3-7) | 2.34 (2-5) | 2.1 (2-3) | 4 (2-11) | 3.1 (2-5) |
| Striatum | 1 | 22.6 (13-28) | 22.4 (10-31) | 4.5 (3-6) | 4.1 (3-8) | 12.2 (9-14) | 12.7 (10-16) | 6.25 (4-8) | 7.8 (5-18) | 2.7 (2-4) | 2.3 (2-4) | 5.3 (3-11) | 4.7 (2-12) |
| | 2 | 22 (13-27) | 22.7 (10-31) | 4.7 (3-8) | 4.7 (3-7) | 12.9 (8-16) | 13.9 (10-17) | 7.4 (4-14) | 6.7 (4-12) | 3.2 (2-9) | 2.7 (2-4) | 6 (3-14) | 5.4 (3-11) |
| Chronic Users | | | | | | | | | | | | | |
| Anterior Cingulate Cortex | 1 | 32.6 (25-46) | 30.4 (18-46) | 2.6 (2-3) | 2.8 (2-4) | 14.5 (9-18) | 13.2 (10-17) | 4.7 (3-14) | 3.7 (2-5) | 2.4 (2-3) | 2.3 (2-3) | 3.9 (2-9) | 4.2 (2-7) |
| | 2 | 33.7 (21-48) | 32 (18-44) | 2.7 (2-4) | 2.9 (2-4) | 13.5 (10-18) | 11.8 (9-15) | 4.1 (3-7) | 3.7 (3-6) | 2.3 (2-3) | 2.2 (2-3) | 3.8 (2-7) | 4.6 (2-7) |

| | | | | | | | | | | | | | |
|---|---|---|---|---|---|---|---|---|---|---|---|---|---|
| Striatum | 1 | 23.8<br>(15-30) | 21.5<br>(14-27) | 3.8<br>(3-5) | 3.75<br>(3-6) | 12.8<br>(9-17) | 13.8<br>(9-19) | 6.5<br>(4-14) | 7<br>(5-13) | 2.4<br>(2-3) | 2.7<br>(2-4) | 5.4<br>(2-12) | 5.7<br>(3-11) |
| | 2 | 23.5<br>(13-32) | 21.1<br>(12-25) | 4<br>(3-6) | 4.2<br>(3-6) | 12.3<br>(11-14) | 11.6<br>(9-15) | 7.9<br>(4-19) | 8<br>(6-13) | 2.5<br>(2-4) | 2.9<br>(2-3) | 5.4<br>(2-10) | 4.8<br>(3-8) |

Table S7. Correlations conducted between absolute average change scores (THC-placebo) of variables that were shown to be significantly affected by THC.

| | Group | Variables | | Correlation | Sig | N |
|---|---|---|---|---|---|---|
| MRS and behavioral | OU | Striatal Glu | SH | -.389 | .211 | 12 |
| | OU | Striatal Glu | NL | .280 | .377 | 12 |
| | OU | Striatal NAAG | SH | -.191 | .211 | 12 |
| | OU | Striatal NAAG | NL | .641 | .025* | 12 |
| | OU | Striatal mI | SH | -.618 | .032* | 12 |
| | OU | Striatal mI | NL | .272 | .392 | 12 |
| | CU | ACC mI | SH | .089 | .794 | 11 |
| FC and behavioral | OU | NAc & VP | SH | -.101 | .755 | 12 |
| | OU | NAc & VP | NL | -.007 | .984 | 12 |

| | | | | | |
|---|---|---|---|---|---|
| OU | NAc & MDN | SH | .065 | .841 | 12 |
| OU | NAc & MDN | NL | -.007 | .984 | 12 |
| OU | VP & MDN | SH | .470 | .123 | 12 |
| OU | VP & MDN | NL | -.224 | .484 | 12 |
| OU | MDN & MC | SH | .182 | .572 | 12 |
| OU | MDN & MC | NL | -.289 | .363 | 12 |
| OU | NAc & MC | SH | -.334 | .289 | 12 |
| OU | NAc & MC | NL | .048 | .882 | 12 |

[*]Significant *P* value

Glu= glutamate; NAAG = total N-acetyl-aspartate; mI= myoinositol; ACC = anterior cingulate cortex; Nac = nucleus accumbens; VP = ventral pallidum; MDN = medial dorsal nucleus; MC = midcingulate area; SH= subjective high; NL= number of lapses on the psychomotor vigilance task, FC= functional connectivity;

Table S8. Significant correlations between functional connectivity and average change scores of the variables of interest.

| Correlation | + / - | Cluster | R or L | BA | k | x | y | z | P value FWE cluster corrected | Pearson's R |
|---|---|---|---|---|---|---|---|---|---|---|
| VAS | + | Middle frontal gyrus | L | 44 | 47 | -44 | 28 | 36 | 0.000 | r=.761, n=12, P =.004 |
| | | Medial frontal gyrus | R | 10 | 19 | 12 | 58 | 10 | 0.004 | r=.708, n=12, P =.010 |
| Number of lapses | + | Middle frontal gyrus | R | 9 | 26 | 34 | 40 | 46 | 0.000 | r=.843, n=12, P =.001 |
| | | Superior frontal gyrus | R | | 49 | 20 | 22 | 48 | 0.000 | r=.641, n=12, P =.025 |
| | | Medial orbitofrontal cortex | L | 11 | 14 | -10 | 34 | -12 | 0.011 | r=.739, n=12, P =.009 |

MNI coordinates of peak voxels for each cluster are given.
